# Watching Others Lift Objects: Corticospinal Excitability is Greater During the Observation of Light than Heavy Lifts

**DOI:** 10.64898/2026.08.27.747509

**Authors:** Orsolya Székely, Janet H Bultitude, Christopher D Chambers, Ezio Preatoni, Jennifer L Davies, Gavin Buckingham

## Abstract

Past studies using transcranial magnetic stimulation have shown larger motor-evoked potentials when people observe someone lifting a heavy object than when they observe someone lifting a light one. This means that observers may engage their own motor system in proportion to the perceived effort. However, the different responses during the observation of light and heavy objects may have been influenced by predictable trial sequences within blocked presentation, making it unclear whether corticospinal excitability reflects online processing of kinematics or is affected by top-down expectations. In this Registered Report, 57 right-handed participants passively observed videos of a precision grip and lift of heavy and light objects while receiving a single-pulse TMS to the left primary motor cortex during the lift phase of the movement. Motor-evoked potentials were recorded from the right first dorsal interosseous muscle. The study compared two main observation contexts: a predictable trial sequence in which repeated videos of the same lifts were presented in a blocked order, and an unpredictable one in which videos were presented semi-randomly and participants could rely only on kinematic cues to perceive the weight of the lifted object. In both conditions, the same videos of lifts of equivalent-looking heavy and light objects were used and only the order of presentation differed. Contrary to our predictions, in the blocked (predictable) condition, there was no significant difference in MEPs elicited by light and heavy lifts. In the unpredictable condition, participants showed greater corticospinal excitability during the observation of the light lifts compared to the heavy lifts. This suggests that in the absence of predictable information, the corticospinal system was sensitive to the observed kinematics, but contrary to previous findings, its excitability varied inversely with the object weight.

## Background

Humans can intuitively interpret others’ actions, recognising information not only in explicit communicative gestures such as a thumbs up, but also in instrumental movements, such as inferring whether an object another person lifts is heavy or light (de C. Hamilton et al., 2007; Runeson & Frykholm, 1981). One explanation behind the latter ability is that observers internally simulate the action they see, engaging their motor system as if performing the action themselves (Gallese et al., 1996; Rizzolatti et al., 1996). To get a quantitative index of motor-system engagement during action observation, corticospinal excitability can be monitored in real time through motor-evoked potentials (MEPs; Barker et al., 1985; Rothwell et al., 1987). Delivering a single pulse of transcranial magnetic stimulation (TMS) to the primary motor cortex (M1) elicits an MEP, the amplitude of which increases with the degree of excitability of the observer’s corticospinal pathway at that moment (Barker et al., 1985; Rothwell et al., 1987).

Early studies on the topic reported greater MEP amplitudes, interpreted as greater motor resonance, when participants observed a video of someone lifting a heavy object compared to a light object (Alaerts et al., 2012; Alaerts, Senot, et al., 2010; Alaerts, Swinnen, et al., 2010). However, almost all of these studies included explicit or implicit predictive cues about the weight of the objects. For example, heavier objects were visibly larger or had a higher filling level, or the lifts were presented in a blocked design (i.e., a block contained only heavy or only light lifts), making the trial order predictable (Alaerts, Senot, et al., 2010; Alaerts, Swinnen, et al., 2010). While the importance of size-based visual cues in making an object’s weight predictable might be obvious, blocked designs create clear expectations as well. In a study by Alaerts et al. (2012), which used blocked presentation, MEPs scaled with the observed object weight not only during the lifting phase but also during the early reach phase, before the object was even touched. At that point, no weight information could be derived from the movement itself, indicating that the predictability of trial order created expectations that influenced motor system responses. Consequently, when visual cues or blocked designs are used, observers can form expectations about the movements, making it difficult to determine whether MEP scaling reflects online processing of kinematic information or is, at least partly, driven by those expectations.

Predictive coding accounts suggest that the motor system continuously generates top-down expectations about others’ actions and updates them when bottom-up sensory information deviates from those predictions (Kilner et al., 2007). From this perspective, making object lifts predictable should strengthen these expectations and potentially amplify motor resonance, whereas removing predictive cues offers a way to test whether motor resonance effects truly reflect the online processing of observed kinematics.

Previous attempts to separate the bottom-up processing of observed kinematics from top-down expectations have yielded mixed results. Alaerts, Swinnen, et al. (2010) used objects whose visible filling level deliberately contradicted their true weight (light objects looked heavy and heavy objects looked light) within a blocked design. MEP amplitudes scaled with the objects’ actual weight, not the misleading visual cue, leading the authors to conclude that kinematic information outweighs expectations. By contrast, Buckingham et al. (2014) used a fully randomised design to test explicit size-based cues. Participants watched videos of an actor lifting two identically weighted but differently sized objects. Larger objects were associated with greater MEP responses, suggesting that observers use their size-based expectations more than the kinematic information. Together, these findings could suggest that when trial order is predictable (blocked design), observers can draw on cumulative kinematic information and discount misleading visual cues, whereas under unpredictable conditions, they rely more heavily on immediate visual appearance.

An apparent exception to this principle is one condition in the study by Senot et al. (2011), who used a fully randomised design without explicit visual cues and still found larger MEPs for heavy lifts in a small sample (n=8). However, this study differs from previous work in key ways. First, it included live action observation (rather than video stimuli) and the actor was aware of the object’s weight. Second, participants interacted with the objects before the experiment started, which may have influenced motor resonance. Moreover, MEPs were analysed using area under the curve (AUC), rather than peak-to-peak amplitude as in most previous studies (Alaerts et al., 2012; Alaerts, Senot, et al., 2010; Alaerts, Swinnen, et al., 2010). These methodological differences do not explain the findings on their own, but limit the ability to make direct comparisons with prior blocked-design studies.

## Objectives

The methodological or design choices of previous studies make it difficult to determine whether weight-dependent MEP modulation reflects real-time processing of lift kinematics or motor preparation driven by expectations. To address this, the present study investigated the effect of trial predictability, presenting lifts either in blocked sequences (all heavy or all light within a block) or in an unpredictable (semi-random, counterbalanced) sequence. As a positive control, we also tested the effect of object appearance, comparing responses to observing lifts of large heavy objects versus small light objects in a condition where the object size indicated weight. The aim of recording peak-to-peak MEPs across these conditions was to show whether corticospinal excitability scales with object weight only when observers can form expectations from order and/or appearance, or whether it also corresponds to the weight of the object when observers must rely solely on bottom-up kinematic cues. The observed pattern of responses was used to provide a measure of motor resonance during action observation. Here, we define action observation as watching (in this case pre-recorded) movements being performed, and motor resonance as the internal representation of these observed motor acts without overt movement, measured via corticospinal excitability.

## Hypotheses

Based on previous research, we had the following hypotheses. First, observing a heavy object would elicit greater MEP peak-to-peak amplitudes than a light object in a blocked (*predictable*) design (H1), consistent with earlier findings. In this condition, we believed that both bottom-up kinematic cues and top-down expectations may contribute to any observed modulation, since repeated exposure to the same weight could have allowed participants to anticipate the object’s properties while still observing movement kinematics. Second, we expected that observing a heavy object would elicit greater MEP amplitudes than observing a light object in an *unpredictable* design (H2). If corticospinal excitability reflects online kinematic processing independent of top-down expectations, we would expect greater MEP amplitudes for heavy lifts even in an unpredictable design. However, if prior knowledge or expectations are important/necessary for this modulation, the effect may be weaker or absent. Third, we expected that the difference in MEP amplitudes between heavy and light objects would be larger in the *blocked* design than in the *unpredictable* design (H3). If prior expectations amplify MEP modulation, the difference in MEP amplitudes between heavy and light objects should be larger in the blocked condition than in the unpredictable condition. For a positive control, we included an additional fourth hypothesis where we predicted that, in an unpredictable design, MEP amplitudes would be greater for heavy objects than light objects when visual cues about objects’ weight were available (H4). We reasoned that in this case, observers could infer weight from object appearance even without predictability from trial order, which would serve as a positive control because if observers do not show greater corticospinal excitability for visually distinct heavy versus light objects, then any absence of effects in the other conditions cannot be confidently attributed to trial predictability. This condition was therefore included to test if our paradigm was sensitive enough to detect weight-related modulation when clear cues were present.

## Methods

### Participants

This study used a within-subjects, repeated-measures design in a single experimental session in counterbalanced order with 57 participants. The mean age of participants was 25.5 years (SD = 9.75, range: 18 to 58). The distribution of biological sex was 33 female (57.89%) and 24 male (42.11%). All participants self-reported to be right-handed. Lastly, 34 (59.65%) of the participants indicated themselves as White, 12 (21.05%) as South Asian, five (8.77%) as East Asian, five (8.77%) as having mixed heritage, and one (1.75%) reported having ‘other’ ethnic background. Participants’ height and body mass were not recorded.

Participants were not screened for history of repetitive movements, skills, or expertise and were not asked to avoid exercise or movement before the experimental session. To control for potential confounds related to temporary physiological fluctuations, participants were asked to refrain from alcohol consumption for 24 hours prior to testing. They were also instructed to maintain their usual sleep patterns and to consume no more or less than their typical amount of caffeine on the day of the experiment. On the day of the experiment, all included participants confirmed full adherence to these instructions. All participants were screened for factors that could influence corticospinal excitability or reaction to TMS:

### Inclusion criteria

- 18 years of age or older
- Right-handed according to self-report

### Exclusion criteria

- History of adverse reactions to TMS, transcranial direct current stimulation, or other forms of brain stimulation
- Personal or family history of seizures, whether epileptic or non-epileptic
- History of stroke
- Previous serious head injury, including neurosurgery or hospitalisation following head trauma
- Presence of metal in the head (outside the mouth), including shrapnel, surgical clips, or welding fragment
- Implanted medical devices such as pacemakers, aneurysm clips, cochlear implants, medical pumps, deep brain stimulators, or intracardiac lines
- Frequent or severe headaches, including a history of recurring migraines
- Current use of, or past use associated with current withdrawal symptoms from, psychiatric or neuroactive medication (e.g., antidepressants, CNS-active drugs such as anticonvulsants), or a history of ongoing and regular non-medical use of psychoactive substances.
- Pregnancy or the possibility of being pregnant
- Dermatological conditions, including eczema or acute skin conditions on the hand that could interfere with electrode placement
- Auditory conditions, such as noise-induced hearing loss, hyperacusis, or tinnitus
- Allergy to adhesive bandages or plasters
- Holding a professional driving licence for heavy goods vehicle, bus, or aircraft
- Any medical conditions that may affect motor function (e.g., Tourette’s syndrome, Parkinson’s disease)

All exclusion criteria are based on the University of Bath TMS safety screening questionnaire (*Supplementary Material 1;* https://osf.io/j3dp7), which was adapted from the one used in the Cardiff University Brain Research Imaging Centre (available as supplementary information in the study by Maizey et al., 2013) and adheres to the safety recommendations outlined by Rossi et al. (2021). The exclusion criteria also included three additional non-TMS-related study specific criteria for participants to have no ongoing (chronic) recurrent or recent (acute) pain in any part of their body, e.g., twisted ankle, or joint replacement (*Supplementary Material 1*). Participants were not screened for further medical, neurological, psychiatric, behavioural, or developmental conditions. There were no tests administered to assess participants’ cognition, but all participants were alert and understood instructions.

The study received ethical approval from the University of Bath, Biomedical Sciences Research Ethics Committee (Reference No. 5817-7604).

### Materials

Participants completed a passive weight-lifting action observation task in which they watched video clips of a male actor, using his right hand, lifting 3-D-printed cylinders that varied in weight and, in one condition, also varied in size. Two cylinders were large and visually identical (height: 7.6 cm, diameter: 10 cm, volume: 596.90 cm^3) but differed in mass: a 200 g (light) and a 1800 g (heavy). A third, smaller cylinder (same height of 7.6 cm with a 7.7 cm diameter and a volume of 353.90 cm^3) also weighed 200 g, providing a visually distinct light object for the cued condition. Comparing this small-light cylinder with the large-heavy cylinder yields a positive-control condition in which weight was perceptually cued. See Figure 1 for a snapshot of each video used.

**Figure 1.**
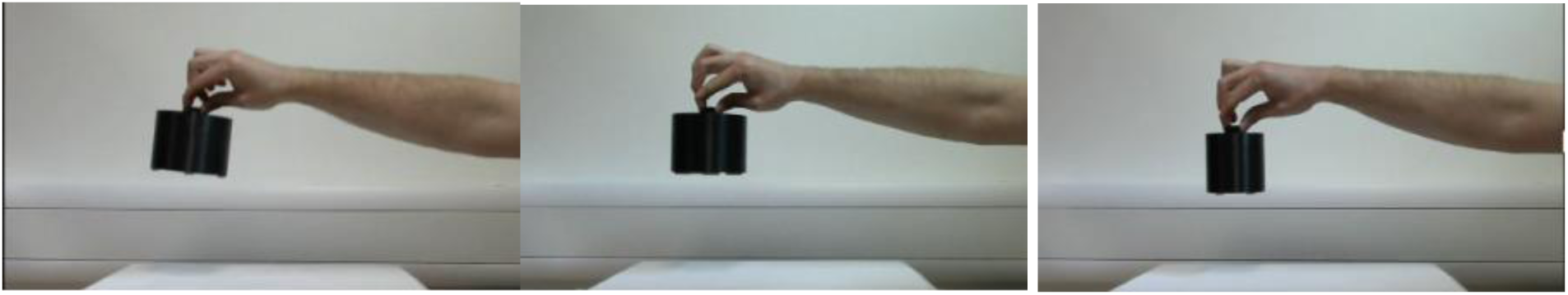
A snapshot from each video. Note that the ‘heavy’ lift 1800g (leftmost) was the same in all three conditions. The large-but-light lift 200g (middle) was the same in the unpredictable and blocked conditions. The smaller light object (rightmost; also 200g) which had the same height and 59.29% of the external volume of the large object, was used in the cued condition only.

All videos were pre-recorded from a third-person, side-on perspective and are provided in *Supplementary Material 2 (*https://osf.io/tn45a). The actor whose hand was featured in the videos was a white male in their mid-20s and performed the lifts with natural kinematics, as the true object weights were unknown to them during recording. Videos were edited so that all trials ended at the same point in the lift and had identical durations. The movement itself was otherwise left unedited. The stimuli were presented on a computer screen about 60 cm from the participant, with PsychoPy (Peirce et al., 2019). Each video lasted 3.54 seconds, and a single TMS pulse was delivered on every trial, randomly timed within the first 0.4 seconds of the lift for the unpredictable and blocked conditions, and within the first 0.5 seconds for the positive control, cued condition. Note, that this is a protocol deviation as our original window for stimulation was 0 - 0.5 seconds for all conditions. Between successive videos, a fixation point was displayed for a random interval of 2.5–3 seconds. This ensured that the timing of the TMS pulse was unpredictable and that there was a gap of at least 5 seconds between pulses, sufficient for preventing carry-over effects from one trial to the next. This stimulation window covers the period in which action observation is expected to modulate corticospinal excitability (Naish et al., 2014) and during which activity in the right first dorsal interosseous (FDI) muscle, which is most directly involved in precision grip, scales with grip and lift force. The combination of variable stimulation timing within the stimulation window and varying intervals between trials ensured that participants could not form expectations about stimulation onset, which could have otherwise influenced corticospinal excitability independently of movement observation. We chose the FDI as our measurement site because it has been suggested to be strongly engaged in the observed action and has been used in similar studies that have shown weight-related MEP differences during the observation of a pinch-grip (Alaerts et al 2010, Senot et al 2011). Our aim was to measure corticospinal modulation by recording from the muscle with the highest expected effect; we did not intend to investigate or make conclusions about either temporal aspects or muscle specificity.

### TMS

TMS was delivered using a Magstim 200² monophasic stimulator (Magstim, Whitland, UK) with a figure-of-eight coil (70-mm diameter per loop, part number: 4150-00; Magstim, Whitland, UK). Coil positioning was continuously monitored using the Brainsight neuronavigation system and software (Rogue Research Inc., Montreal, Canada). TMS pulses were externally triggered via Psychopy (Peirce et al., 2019) using embedded Python code. The stimulator was connected to the stimulus presentation computer via a USB-to-serial adapter and was controlled using the MagPy package in Python (McNair, 2017). The full Psychopy task, including embedded Python code for TMS control, is available in *Supplementary Material 3 (*https://osf.io/72fmu).

### EMG

Interference EMG was recorded using the Brainsight NIBS EMG kit (Rogue Research Inc., Montreal, Canada). This system includes the Model 3 amplifier (overall gain 4 444 V/V, bandwidth 16 to 470 Hz) integrated with the Brainsight acquisition software. After cleaning the skin with alcohol wipes, surface Ag/AgCl electrodes (pre-gelled disposable; Kendall™ 130) were placed over the relaxed right FDI muscle in a tendon–belly monopolar montage, with the active electrode positioned over the muscle belly, the reference electrode placed on the proximal phalanx of the index finger, and a ground electrode positioned over the ulnar styloid (wrist bone). Signals were digitised at a 3 kHz sampling rate.

### Procedure

Experimental sessions took place at varying times between 9 am and 6 pm. Excluding arrival, consent, demographic questionnaires, and safety screening, the experimental session, including set-up, lasted a mean of 92.8 minutes (SD= 8.4; range: 77 to 120). When signing up to participate, participants received the TMS safety-screening form (with the items mentioned under Exclusion Criteria) by e-mail so that they could review the questions and, if necessary, withdraw. On arrival, they gave written informed consent, completed the safety screening form, and filled in a brief demographic questionnaire (age, sex, ethnicity, handedness). Following consent and screening, participants were seated comfortably in an armchair and were instructed to remain at rest (and stationary) throughout the experiment (with the possibility to move around after set-up and after every 11 trials during the experiment). All participants wore earplugs during TMS to attenuate the effect of stimulation sound. To familiarise them with the stimulation, a short series of very low-intensity pulses were delivered, starting at about 25% of maximum stimulator output (MSO) and gradually increasing to 50 to 60% MSO, which was confidently and visibly above each participant’s rMT, making sure they would be comfortable with suprathreshold stimulation during the experiment. After the familiarisation, the EMG setup was prepared and, using real-time Brainsight feedback, the experimenter identified the FDI motor hotspot, defined as the scalp position where single-pulse TMS evoked the largest and most consistent MEPs in the contralateral (right) relaxed FDI muscle. At the hotspot, resting motor threshold (rMT) was defined as the lowest intensity that elicits MEPs of ≥ 50 µV peak to peak in 5 of 10 consecutive trials (Rothwell et al., 1999). Throughout this setup and the entire experiment, the coil was held at a 45° angle to the sagittal midline of the head to induce a posterior–anterior current in the underlying neural tissue, with its position monitored in Brainsight.

Before beginning the first block, participants completed a short practice session, consisting of watching five video trials without TMS and then five with TMS to ensure they were comfortable with the procedure. Stimulation was set to 130% of the individual’s rMT and to the previously defined FDI motor hotspot. This intensity level was selected to replicate the methodology of studies by Alaerts and colleagues (Alaerts et al., 2010a; Alaerts et al., 2010b; Alaerts et al., 2012; Senot et al., 2011), which consistently applied 130% rMT. During the experiment, participants rested their right hand with the EMG electrodes on their lap or on the table, whichever they found more comfortable and yielded the least noisy EMG signal. To replicate previous experiments (Alaerts et al., 2010a; 2010b; 2012) as closely as possible, the instructions to participants were to simply observe the lifting actions while remaining attentive. Instructions were delivered as spoken words and were simultaneously displayed on the screen in front of participants (see *Supplementary Material 4* for the script of these instructions; https://osf.io/u35sn). As in the studies by Alaerts and colleagues (Alaerts et al., 2010a; Alaerts et al., 2010b; Alaerts et al., 2012), adherence to instructions was not explicitly monitored; however, after every 11 trials, there were short breaks provided, and participants were observed by the experimenter throughout to make sure they were alert. Within the breaks, they were able to stand up or stretch while keeping the electrodes in place.

The experiment consisted of three counter-balanced blocks of 44 trials each (22 heavy, 22 light in each block). In the unpredictable block, the two objects were visually identical and appeared in a prespecified pseudorandom order with no more than three identical trials in succession. In the predictable block, the same identical-looking weights were presented in two mini-blocks of only heavy and only light objects, with the presentation order counterbalanced between participants. In the cued unpredictable condition, the weights were visually distinct (small = light, large = heavy) and followed the same semi-random order as the unpredictable condition. There were 132 TMS pulses delivered in total per participant during the experimental task.

During each trial, participants watched a video of a hand lifting an object, and a TMS pulse was delivered at a random time within a 0.4-second (blocked and unpredictable conditions) and 0.5-second (cued condition) window corresponding to the transition from the late grasp-to-lift phase into the lift itself. In this phase, the lifter, who did not know the object’s weight in advance, must scale their grip and load forces to the actual weight in order to lift it, making this the time window most likely to reveal weight-specific kinematic information. To verify that the randomised stimulation window was comparable across conditions, we computed the mean pulse time (relative to lift onset) for each condition and assessed whether these differ (t-test comparing heavy vs light trials within each condition).

At the end of the TMS blocks, participants were asked how many different object weights they believed they observed to gain insight into whether they were consciously aware of the categorical differences in object weights. After the three TMS blocks, participants completed two brief behavioural blocks (weight judgement task) of 20 trials each (40 in total) without TMS, using the same video stimuli as before. One block included 20 lifts of the visually identical light and heavy objects, presented in an unpredictable order. The other block included the visually distinct weights (10 light and 10 heavy) presented in an unpredictable order. The order of the blocks was counterbalanced. After each video, participants rated the perceived weight of the object by a mouse click on a continuous scale anchored from “lightest” to “heaviest”. These ratings informed the understanding of whether any differences in MEPs corresponded to conscious differentiation between these objects. See *Supplementary Material 5* for the PsychoPy script of this weight-judgement task, adapted from the task used in our previous study (Székely et al., 2026); https://osf.io/5pebh. See *Figure 2* for a schematic representation of the experimental procedure. At the end of the experiment, participants were asked whether they had previously taken part in action observation research.

**Figure 2.**
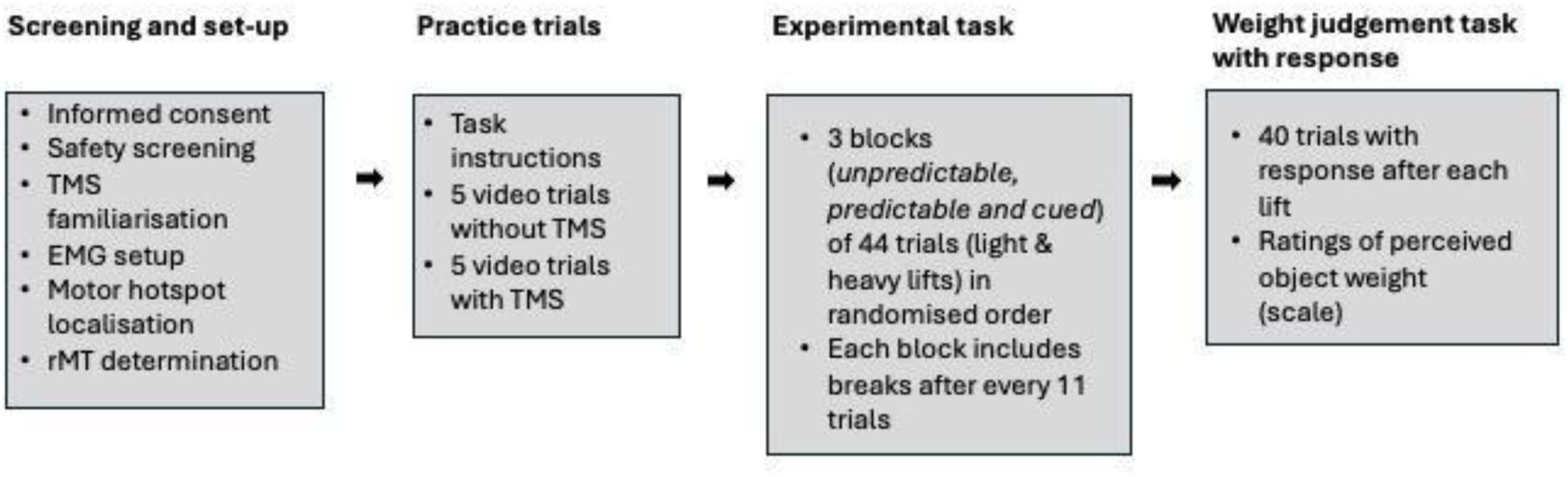
Overview of experimental procedure.

### Data Analysis

All data-analysis codes were written in R and are available on the Open Science Framework. (*Supplementary Material 6;* https://osf.io/nykfp/files/d8qkb). MEP amplitude was used as the dependent variable for all hypotheses tested, defined as the maximum minus minimum EMG peak within the window of 10-50 ms after each TMS pulse. MEP data were normalised to the maximum MEP across all conditions and trials (within each participant) to account for different factors such as electrode placement, muscle size, threshold, and MEP size, that likely vary between participants and that were not of interest within our study.

Background EMG was quantified in the 100 ms preceding stimulus onset. A trial was discarded if the background EMG exceeded 50 µV peak-to-peak. Trials in which coil position deviated by more than 2 mm from the motor-hotspot target were excluded. Data from participants who did not have at least 15 usable MEP trials per condition and weight after these filtering steps were excluded (minimum trial number required for the reliable assessment of corticospinal excitability in our sample based on the method described by Biabani et al., 2018 [*Supplementary Material 7*; https://osf.io/nykfp/files/wqs6x]). Note that this is a protocol deviation as our original criteria for the minimum number of usable trials was 20. Participants were excluded and replaced if they withdrew before the end of the TMS blocks. They were also excluded if a reliable motor hotspot or rMT could not be identified (reliability assessed based on the subjective judgement of the experimenter), in which case, the experiment did not start. For details of the statistical analysis for each hypothesis, see *Table 1*.

**Table 1.** Study Design Table.

| Question | Hypothesis | Sampling plan | Analysis Plan | Rationale for deciding the sensitivity of the test for confirming or disconfirming the hypothesis | Interpretation given different outcomes | Theory that could be shown wrong by the outcomes | Results |
| --- | --- | --- | --- | --- | --- | --- | --- |
| Q1: Does observing a heavy vs. light lift modulate corticospinal excitability in a blocked design? | H1: MEP amplitudes will be significantly larger for heavy vs. light lifts in the blocked condition | Recruiting a minimum of 40 participants, aiming for 53 participants, final sample: 57 participants | Paired-samples t-test on MEP peak-to-peak amplitudes for heavy vs. light trials in the blocked condition | See the *Effect size estimation* section for the details of sample size estimation. | *Significant heavy > light:* confirms that repeated predictable observation elicits weight-dependent MEP scaling.<br>*No difference or light > heavy:* the hypothesis that repeated predictable observation elicits weight- dependent MEP scaling is **not** confirmed. | Predictable repetition allows observers to integrate kinematic cues and generate weight-specific corticospinal facilitation. | MEP amplitudes were not significantly different for heavy vs. light lifts in the blocked condition. |
| Q2: Does observing a heavy vs. light lift modulate corticospinal excitability in an unpredictable design? | H2: MEP amplitudes will be significantly larger for heavy vs. light lifts in the unpredictable condition. |  | Paired-samples t-test on MEP peak-to-peak amplitudes for heavy vs. light trials in the unpredictable condition |  | Significant heavy > light: supports the idea that bottom-up kinematics alone can drive MEP modulation.<br>*No difference or light > heavy: the hypothesis that bottom-up kinematics alone can drive MEP modulation is **not** confirmed.* | Online kinematic information is enough to modulate corticospinal excitability in the absence of explicit or implicit cues. | MEP amplitudes were significantly larger for light vs. heavy lifts in the unpredictable condition. |
| Q3: Is the heavy–light difference in | H3: The MEP difference |  | Paired-samples t-test |  | *Interaction significant: shows prior* | Top-down prediction enhances MEP | The MEP difference between heavy |
| MEPs larger in blocked vs. unpredictable designs? | between heavy vs. light lifts will be significantly larger in blocked than in the unpredictable condition. |  | comparing the difference of the difference between the light and heavy weights of the blocked and unpredictable conditions. |  | expectation amplifies the heavy-light MEP contrast. *No interaction:* the hypothesis that prior expectation amplifies the heavy-light MEP contrast is **not** confirmed. | beyond that driven by kinematics alone. | vs. light lifts was not significantly different in the blocked than in the unpredictable condition. |
| Q4: Does observing a heavy vs. light lift modulate corticospinal excitability in an unpredictable but cued (visually distinct) design? | H4: MEP amplitudes will be significantly larger for heavy vs. light lifts in the cued (visually distinct) condition. |  | Paired-samples t-test on MEP peak-to-peak amplitudes for heavy vs. light trials in the cued (visually distinct) condition |  | *Significant heavy > light:* demonstrates that explicit size cues can elicit weight-specific facilitation even without predictable order. | That kinematic information and visual cues to object weight are together sufficient to drive weight-related changes in corticospinal excitability, even when trial order is not predictable | MEP amplitudes were not significantly different for heavy vs. light lifts in the cued (visually distinct) condition. |
|  |  |  |  |  | *No difference*<br>*or*<br>*light > heavy:*<br>the hypothesis that explicit size cues can elicit weight-specific facilitation even without predictable order is **not** confirmed. |  |  |

*Note.* All preregistered, and further exploratory hypotheses were tested with $\alpha = 0.02$ independently of each other. The analyses reported in this table deviate from the registered Stage 1 protocol. These deviations are described in detail in the Deviations from the Preregistered Stage 1 Protocol section below. Any other analyses below, not mentioned here, are interpreted in a post-hoc manner as exploratory findings. We collected additional data for exploratory analysis, such as MEPs from the abductor pollicis brevis muscle (APB), another muscle involved in precision grip. We also analysed additional variables such as area-under-the-curve to provide further context for interpreting the study's findings. These measures fell outside the scope of the Registered Report, and we did not have sufficiently specific predictions or the statistical sensitivity to design the study around them.

### Pilot data

We collected pilot data to assess the feasibility of the experiment. All four pilot participants found the task manageable. The pilot tested the blocked presentation of two visually identical but different weighting objects. This configuration was expected to produce larger effects than the unpredictable presentation, but smaller than the size-cued presentation. The pilot data showed a large within-subject effect (dz = 1.05; see Figure 3). The data are visualised in Figure 3. The complete pilot dataset and the accompanying R analysis code are available in *Supplementary Materials 8 and 9 (see* https://osf.io/25hvp *and* https://osf.io/ua2n9). Pilot participants did not complete the full experimental session, only the condition corresponding to H1. The data collected were not reused in the confirmatory preregistered analysis.

**Figure 3.**
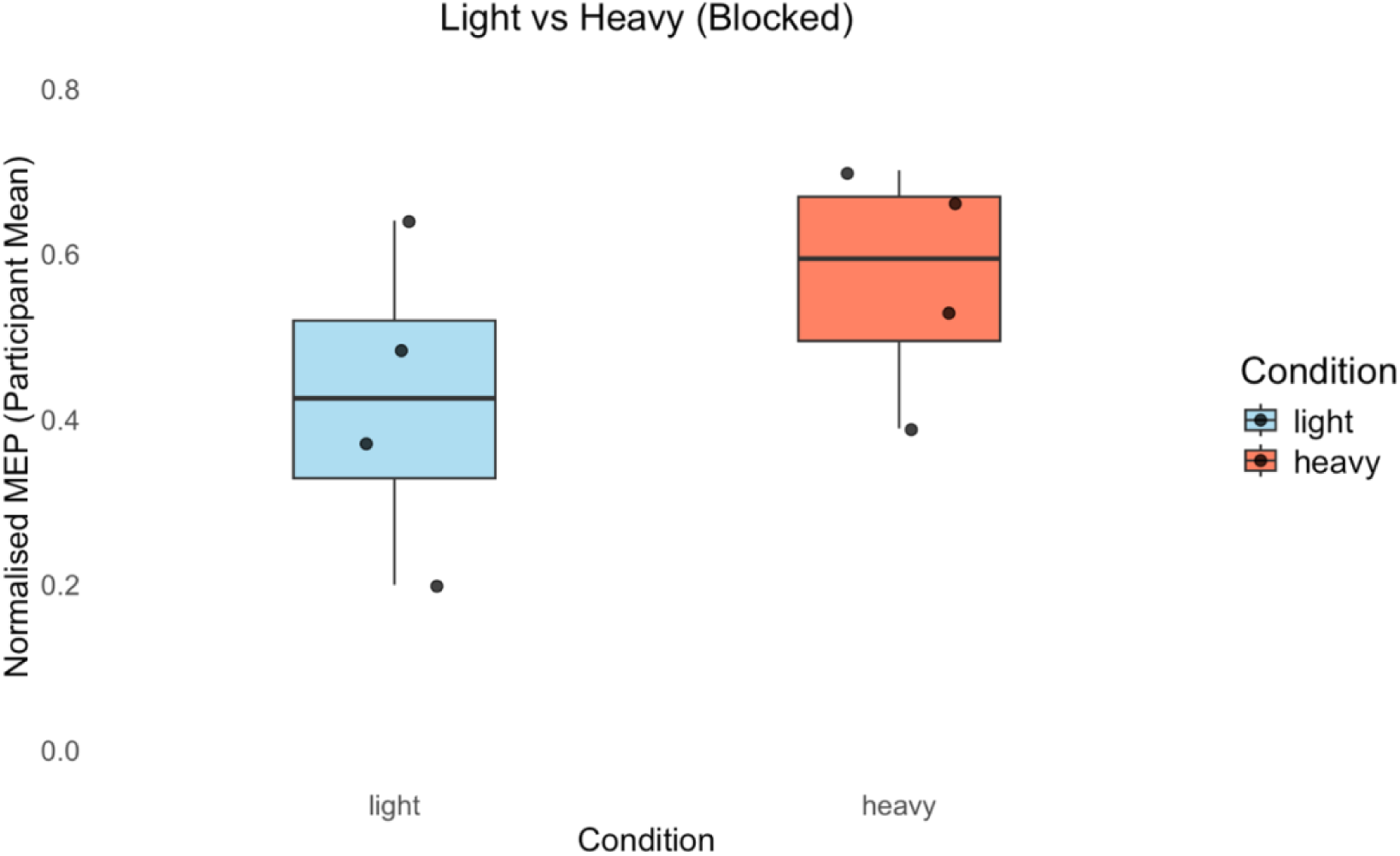
Mean Normalised MEP amplitude in the Blocked Condition with two weights from 20 trials per person during the piloting of the experiment. The boxes represent the interquartile range (IQR). The horizontal line inside the box indicates the median.

### Effect Size Estimation

See the four hypotheses and details in *Table 1.* To establish the smallest estimated effect size of interest, we used the effect from the study by Senot et al (2011) from their ‘hidden’ condition (i.e., identical-looking weights presented in an unpredictable, random order, n=8). As the data required to compute an effect size were not reported, we digitised Fig. 2B using a web-based plot digitiser, which yielded an estimated effect size of dz=1.01. This effect was considered as the smallest estimated effect size of interest, as it comes from an unpredictable, uncued design within that study, which corresponds to the condition in our study where we expected the smallest effect. In our other conditions, where predictive cues were available, either visual or based on trial presentation, we expect larger effects. H2 within our study repeated this exact, un-cued, unpredictable condition, so this effect directly reflects an estimate of the heavy-light difference in that condition. H1 (predictable order) and H4 (size-cued positive control) address the same heavy-light MEP contrast in conditions where the true effect is estimated to be larger than the uncued unpredictable benchmark. For H3, we tested whether the MEP difference between heavy and light weights was larger in the blocked design than in the randomised design. This analysis concentrated on the difference of a difference, meaning the effect size was expected to be smaller than for H1, H2 or H4. As a heuristic, we used the lower bound of the 80% confidence interval of the effect size from the study by Senot et al. to estimate a plausible minimum effect size that we considered meaningful to detect for this hypothesis. The lower bound of the 80% confidence interval for Cohen’s dz effect size from the study by Senot et al. is dz=0.52. Based on the variability across participants in our pilot data (SD of the heavy-light differences), an effect size of dz = 0.52 would correspond to a standardised MEP difference of 0.073. Effects smaller than this threshold would have limited theoretical relevance, and therefore, we did not power the study to detect effects below this level.

We used the pwr package in R (*Champely, 2020)* to perform the power analyses. The R code used for effect size estimation is available in *Supplementary Material 10 (*https://osf.io/mzsq4). For each hypothesis, the primary statistical contrast was identified to be tested in the analysis (e.g., the heavy–light main effect for H1, H2, and H4; the Weight × Design interaction for H3). We then conducted the power analysis as a single one-degree-of-freedom comparison (equivalent to a paired-samples t-test for within-subject contrasts or the relevant one-degree-of-freedom term from the linear model). This ensured that the power analysis was identical in form to the confirmatory statistical tests that were applied to each hypothesis, while avoiding inflation from model-specific complexity.

An estimated realistic minimum target sample we committed to due to time and resource limitations was n=40. With α = 0.02 and 90 % power, this sample size would have been sufficient to detect the effect size of dz = 0.59, which is over 40% smaller than both the effect size estimated from the Senot et al. study (dz = 1.01) and the effect size observed in our pilot data (dz = 1.05). On this basis, n=40 was considered as suitable to detect the smallest plausible estimated effect for H1, H2, and H4.

For H3, which includes an interaction effect and a smaller estimated minimum plausible effect of interest (dz = 0.52), we required n = 51 for 90 % power at α = 0.02. We aimed to recruit up to n = 53 participants to ensure this level of sensitivity. However, if recruitment had stopped at n = 40 due to practical constraints, the analyses would have been reported for H1, H2, and H4, and any implications of reduced sensitivity for H3 would have been explicitly addressed in the interpretation of results.

We report the methods in accordance with reporting guidelines for action observation and TMS studies, including the Guidelines for Reporting Action Simulation Studies (GRASS; Moreno-Verdú et al., 2024; see *Supplementary Material 11* for the filled checklist; https://osf.io/nykfp/files/gkurj) and the Transcranial Magnetic Stimulation Reporting Assessment Tool (TMS-RAT; https://tms-rat.org/; Székely et al., 2026).

### Deviations from the Preregistered Stage 1 Protocol Major deviations

After completing data collection, we discovered an error in the stimulation code, and made two changes to the protocol to manage some resulting irregularities in our stimulation timings. The error was discovered after the initial inspection of the results, and the deviations were finalised based on theoretical bases, without any assessment of the consequence of their effect. First, we use a consistent stimulation timing window of 0-0.4 seconds (instead of 0-0.5 seconds) across the blocked and unpredictable conditions for all participants to make these two conditions completely comparable. The cued condition, which was included as a positive control and was not directly compared with the blocked or unpredictable conditions in any of the preregistered analyses, retained the 0-0.5 second stimulation window for both light and heavy trials.

Second, due to the need to exclude a small number of trials in the blocked and unpredictable conditions with stimulation timing of greater than 0.4, we now use a minimum of 15 valid trials per condition and weight (instead of the 20 that was described in the stage 1 protocol; see *Supplementary Material 7* for the theoretical argument for why this is sufficient; https://osf.io/nykfp/files/wqs6x). To confirm that these deviations do not affect the conclusions, we conducted sensitivity analyses using alternative minimum-trial thresholds and the originally collected stimulation times. These checks are reported in the Results and detailed in the corresponding Supplementary Materials.

### Minor deviations

In the Stage 1 protocol, the planned analyses for H1, H2, and H4 were presented in *Table 1* using linear models of the form MEP ∼ Weight. However, the corresponding power analyses were conducted as one-degree-of-freedom within-subject contrasts, which is equivalent to paired-samples t-tests on heavy–light differences. Because the hypotheses concern within-participant differences between two weight conditions, and because treating trial-level observations as independent would be incorrect, the confirmatory tests were conducted as paired-samples t-tests on participant-level condition means.

Timing logs were not recorded for blocked-condition trials in participants who saw light-object trials first. Therefore, trial-level stimulation timing could not be verified for these trials in the same way as for the other conditions. The available timing logs from the other conditions showed that pulses were delivered as expected from the stimulation code.

For two participants, the available pre-pulse baseline EMG window was 50 ms rather than the preregistered 100 ms because 50 ms was the default setting in the Brainsight system, and this setting was not changed during the experimental set-up. These participants were retained, and the available 50-ms pre-pulse window was used for baseline EMG screening.

The final included sample was 57 participants. This exceeded the originally planned target of 53 participants because the revised trial-inclusion criterion allowed more participants to be retained after trial-level exclusions.

The Stage 1 protocol stated a lower age limit of 16 years. However, ethical approval was granted to collect data from participants aged 18 or older, so we used a lower age limit of 18 years.

## Results

### Data exclusions

The raw EMG data, demographic and motor characteristics, and behavioural weight judgement task data are available in *Supplementary Materials 12* (https://osf.io/nykfp/files/x8eah*), 13* (https://osf.io/nykfp/files/rwg9m), and *14* (https://osf.io/nykfp/files/6n7xr), respectively. Complete data were initially available from 65 participants, comprising a total of 8580 TMS trials. Based on the registered exclusion criteria, 271 trials (3.16%) were excluded because the pre-pulse background EMG exceeded 50 µV in the FDI muscle due to participant movement or noise. A further 31 trials (0.36%) were excluded because the coil position deviated more than 2 mm from the target (motor hotspot for FDI). Then, another 253 (2.95%) trials were excluded because the stimulation timing was outside of the 0.4 second stimulation window. Finally, eight participants (12.31%) were excluded because they had fewer than 15 trials after these exclusions for at least one condition per weight, leaving 57 participants, with a total of 7186 trials. In the final included trials, the mean coil-position deviation from the FDI motor-hotspot target was 0.50 mm (SD = 0.29, range: 0.02 to 1.99 mm).

### Motor characteristics of participants

Only six (10.53%) out of the 57 participants reported previously having taken part in action observation related research. The mean rMT of participants was 41.80% of MSO (SD=7.12, range: 22.50 to 55.50 with interpolation when needed). As stimulation intensity was set to 130% of resting motor threshold and rounded to the nearest whole percentage of MSO, the average stimulation intensity during the experiment was 129.93% rMT (SD = 0.73, range: 128.89 to 131.43) and 54.13 % of MSO (SD = 9.33, range: 29 to 72).

### Perception of object weights

At the end of the TMS blocks, before the behavioural (weight judgement) task, participants reported how many different object weights they believed they observed during the TMS experiment. It was specified that this refers to the number of weights, and not the number of videos or object sizes. The mean perceived number of weights was 4.27 (SD=2.54, range 2 to 15). Only five participants (8.77%) reported correctly that they saw only two different weighted objects.

The behavioural weight judgement data were available for 54 of the 57 participants (data from three participants were not available due to technical/procedural interruptions during data collection). Ratings were scaled to a 0-100 scale, where 0 indicates the lightest object and 100 indicates the heaviest object. Participants reliably differentiated between the light and heavy weights. In the unpredictable condition, heavy objects were perceived as heavier (M = 77.8, SD = 11.1) than light ones (M = 20.0, SD = 14.4), mean difference = 57.8, t(53) = 24.98, p < .001. In the cued condition, the ratings were also higher for heavy objects (M = 80.6, SD = 11.5) than for light objects (M = 18.8, SD = 9.43), mean difference = 61.7, t(53) = 29.43, p < .001. See Figure 4 for the visualisation of the results of the weight judgement trials.

**Figure 4.**
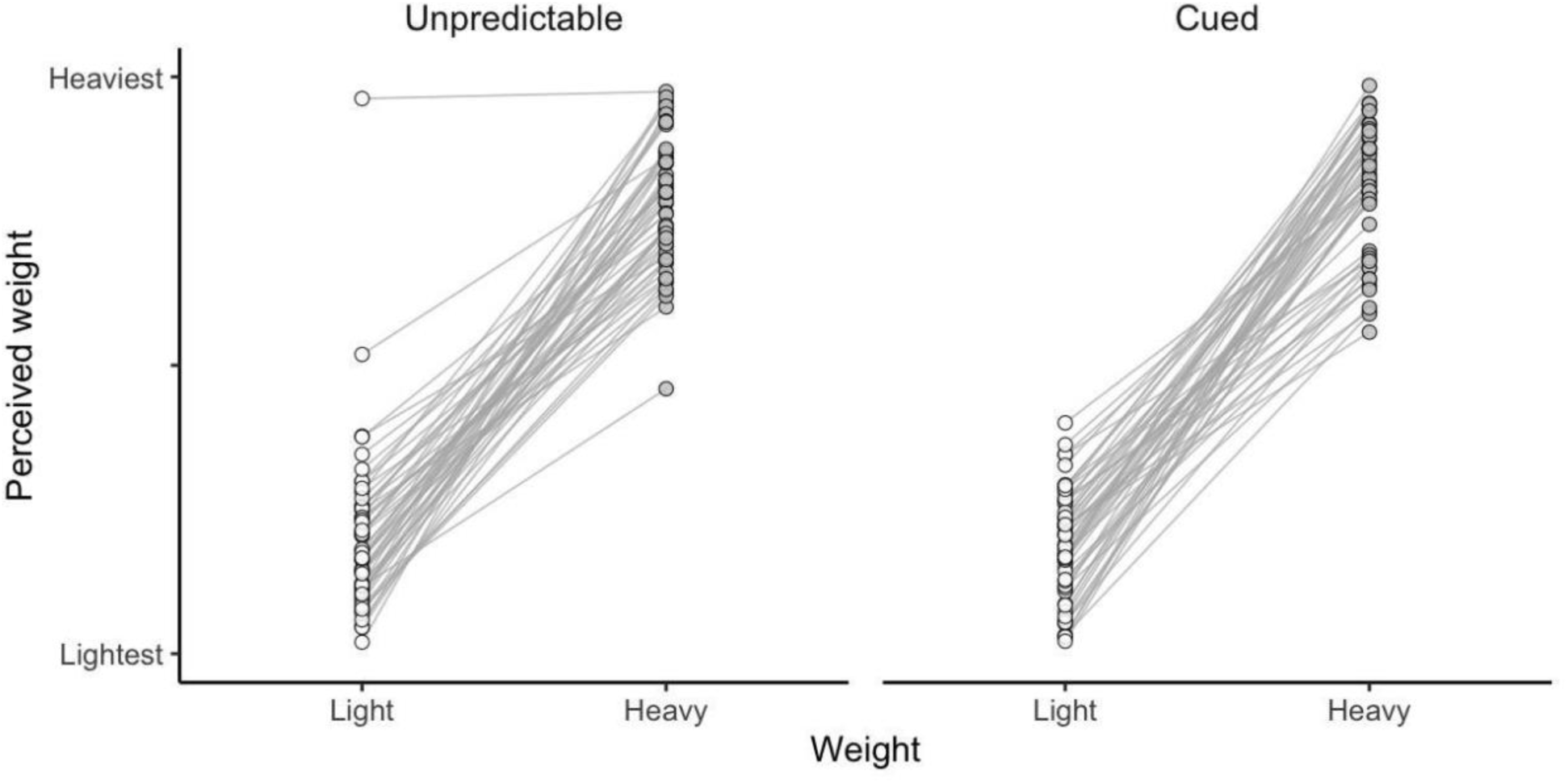
Mean responses from each participant (n=54) for each weight over 20 unpredictable (10 light and 10 heavy) and 20 cued (10 light and 10 heavy) trials of the behavioural weight judgement task. ‘Lightest’ corresponded to 0 and ‘Heaviest’ corresponded to 100 on the Y axes, but these numbers were not shown to participants; they only saw an unnumbered scale with the two labels at the extremes.

### H1: MEP amplitudes will be significantly larger when observing heavy vs. light lifts in the blocked (*implicit predictable*) condition

The results are summarised in Figure 5. A paired-sample t-test showed that there was no difference in the normalised MEP amplitude elicited in the FDI muscle during observation of heavier (Mean= 0.40, SD=0.21) versus lighter (Mean= 0.42, SD=0.21) lifts in the blocked condition. The mean response difference between heavy and light lifts was -0.021 (2.1%), t(56) = -1.51, p = 0.13, dz = -0.20. An exploratory Bayesian paired t-test showed that these results were 2.37 times more likely under the null than the alternative hypothesis (BF₁₀ = 0.42; BF₀₁ = 2.37), suggesting inconclusive evidence.

**Figure 5.**
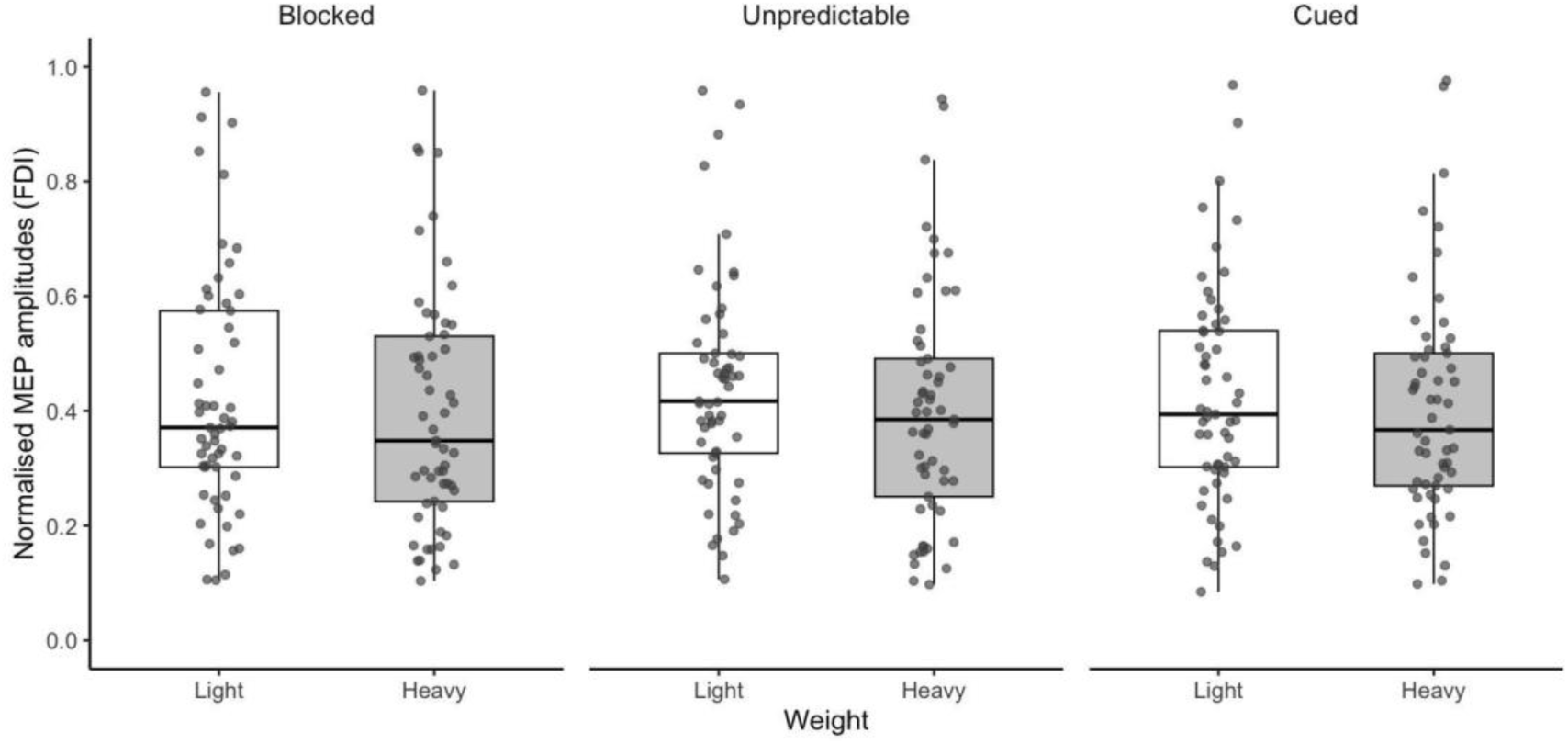
Normalised peak-to-peak MEP amplitudes for light and heavy object lifts in the blocked, unpredictable, and cued conditions in the FDI muscle. Points show participant-level condition means (n=57), the boxes show the interquartile range, whiskers show 1.5 x the interquartile range, and the horizontal lines show medians.

Thirty of the included participants started the blocked condition with heavy trials, and 27 with light trials. To assess whether the order made a difference, we compared the difference between the normalised MEP amplitude during the observation between heavy and light lifts between the two versions of the blocked condition (light first versus heavy first). The overall tendency showed that participants who took part in the heavy trials first had slightly larger difference between the MEPs during the light and heavy lifts (with greater response to the light lifts), but this difference was not significant based on our preregistered significance threshold of <0.02, Welch independent t-test: t(49.73)= -2.22, 95% CI: -0.118 to -0.006, p = 0.031.

### H2: MEP amplitudes will be significantly larger when observing heavy vs. light lifts in the unpredictable condition

In the unpredictable condition, a paired-sample t-test showed that there was a significantly larger normalised MEP elicited in the FDI muscle during observation of light (Mean= 0.44, SD=0.19) compared to heavy (Mean= 0.40, SD=0.20) lifts, with a mean difference of -0.040 (4.0%); t(56) = -4.19, p < 0.001, dz = -0.56 (Figure 5). An exploratory Bayesian paired t-test showed that these results were 223.64 times more likely under the alternative than under the null hypothesis (BF₁₀ = 223.64; BF₀₁ = 0.0045), suggesting very strong evidence for this effect.

### H3: The difference in MEP amplitudes between heavy and light objects will be larger in the *blocked* condition than in the *unpredictable* condition

We compared the difference between the MEP response during heavy and light lifts between the blocked and unpredictable conditions. A paired t-test showed no significant difference of difference, mean difference = 0.018, t(56) = 1.06, p = 0.29, 95% CI = -0.016 to 0.053, dz = 0.14 (Figure 5). An exploratory Bayesian paired t-test indicated that the data were 4.057 times more likely under the null than the alternative hypothesis (BF₁₀ = 0.24; BF₀₁ = 4.057), suggesting moderate evidence for the null hypothesis.

### H4: Observing a heavy object will elicit greater MEP amplitudes than observing a light object in the cued (visually distinct) condition

In the explicit (visually) predictable design, where the heavy and the light objects were different sizes to one another, and were thus visibly different sizes, we found no difference between the response to observing heavy (Mean= 0.41, SD=0.19) vs light (Mean= 0.43, SD=0.19) object lifts (mean difference: -0.021, t(56)= -2.086, p = 0.042, 95% CI= -0.041, 0.00083, dz = -0.28; see Figure 5). An exploratory Bayesian paired t-test showed that these results were only 1.076 times more likely under the null than the alternative hypothesis (BF₁₀ = 0.93; BF₀₁ = 1.076), suggesting inconclusive evidence.

## Exploratory Robustness checks

### Choice of normalisation method and outcome measure

We have performed multiple exploratory analyses to check the robustness of our findings. The details of the results of all statistical analyses are available in *Supplementary Material 15 (*https://osf.io/nykfp/files/hq2t3). In summary, we repeated all analyses addressing the four preregistered hypotheses: 1) without normalising the FDI MEP peak-to-peak amplitude response to assess the effect of the choice of normalisation; and 2) using the normalised AUC instead of peak-to-peak MEP amplitude as outcome, to assess the effect of the choice of outcome measure. The results were unchanged; normalisation and the choice of outcome measure did not change the results of the four pre-registered analyses.

### Balance of stimulation timing within conditions

We compared the mean pulse timing for included trials from the start of the stimulation window between heavy and light trials in each condition. There was no significant difference between pulse timings between heavy-light lifts in the cued condition, t(56) = 1.15, p = 0.26, in the unpredictable condition, t(56) = -1.02, p = 0.31, or in the blocked condition, t(29) = 0.49, p = 0.63. In terms of absolute timing from video onset, the blocked and unpredictable conditions were matched across weights: object contact, and therefore the start of the stimulation window, occurred 3.0 seconds after video onset for both light and heavy lifts. However, in the cued condition, the stimulation window began 2.3 seconds after video onset for the small-light object and 3.0 seconds after video onset for the large-heavy object, because the natural reach toward the smaller object was faster. Consequently, relative to video onset, there was no significant difference between the stimulation window of the heavy and light lifts in the blocked condition, t(29) = 0.49, p = 0.63, or the unpredictable condition, t(56) = -1.02, p = 0.311. However, in the cued condition, pulse timing differed between weights, t(56) = 111.00, p < .001.

## Effect of protocol deviations

### The effect of not applying the new data exclusions that were introduced as protocol deviations

As described under ‘Deviations from the Preregistered Stage 1 Protocol’, we changed the stimulation timing window from 0-0.5s to 0-0.4s in the blocked and unpredictable conditions to make these conditions comparable and reduced the minimum number of trials required for retention of a participant in the analysis from 20 to 15. To assess whether these deviations affected the results, we repeated the preregistered analyses without excluding trials with stimulation timing at 0.5 seconds after object contact from the blocked and unpredictable conditions, and using a minimum of 20 trials per condition and weight as preregistered in Stage 1. When 0.5-second stimulation trials were not excluded, and a minimum of 20 trials per condition and weight were needed, 53 participants were included in the analysis. Although this analysis was imbalanced by the 0.1 second difference in the size of the pulse timing window in the blocked condition (H1) and the comparison of the blocked and the unpredictable conditions (H3), the results for the four preregistered hypotheses were unchanged from those presented above, suggesting that the findings are robust to the impact of the unregistered data exclusions that we applied (see Table 2).

**Table 2.** Results of the analyses without applying the new (unregistered) data exclusions.

| Analyses | N | Mean<br>diff. | 95%<br>CI<br>lower<br>bound | 95%<br>CI<br>upper<br>bound | t-value | df | p-value | dz | BF10 | BF01 |
| --- | --- | --- | --- | --- | --- | --- | --- | --- | --- | --- |
| H1 | 53 | -0.028 | -0.058 | 0.001 | -1.94 | 52 | 0.057 | -0.27 | 0.85 | 1.17 |
| H2 | 53 | -0.042 | -0.062 | -0.022 | -4.22 | 52 | < .001* | -0.58 | 226.53 | 0.004 |
| H3 | 53 | 0.013 | -0.024 | 0.05 | 0.72 | 52 | 0.472 | 0.1 | 0.19 | 5.21 |
| H4 | 53 | -0.018 | -0.039 | 0.003 | -1.69 | 52 | 0.097 | -0.23 | 0.56 | 1.77 |

For the cued condition specifically, we also repeated H4 after restricting the analyses to trials with stimulation within the 0.4 seconds after object contact. The results were unchanged with no significant difference between peak-to-peak normalised MEPs between light and heavy weights, t(52) = -1.62, p = 0.11, dz = -0.22, BF10 = 0.51, BF01 = 1.97.

### Number of minimum required trials per participant

To check the effect of using different minimum numbers of trials per condition and weight, we repeated the analyses for minimums of every number between 10 and 20 (other than 15, which is the minimum number of trials for the main analysis reported above). The number of participants possible to include based on this ranged from 63 to 23 (see Table 3). When we repeated the main analyses using the data for each of these further ten datasets, analyses showed the same results: i.e. the only significant effects were in the unpredictable condition where participants had a larger MEP response to TMS while observing light compared to heavy lifts (see the detailed results of these analyses in *Supplementary Material 16;* https://osf.io/nykfp/files/9nbcx).

**Table 3.**
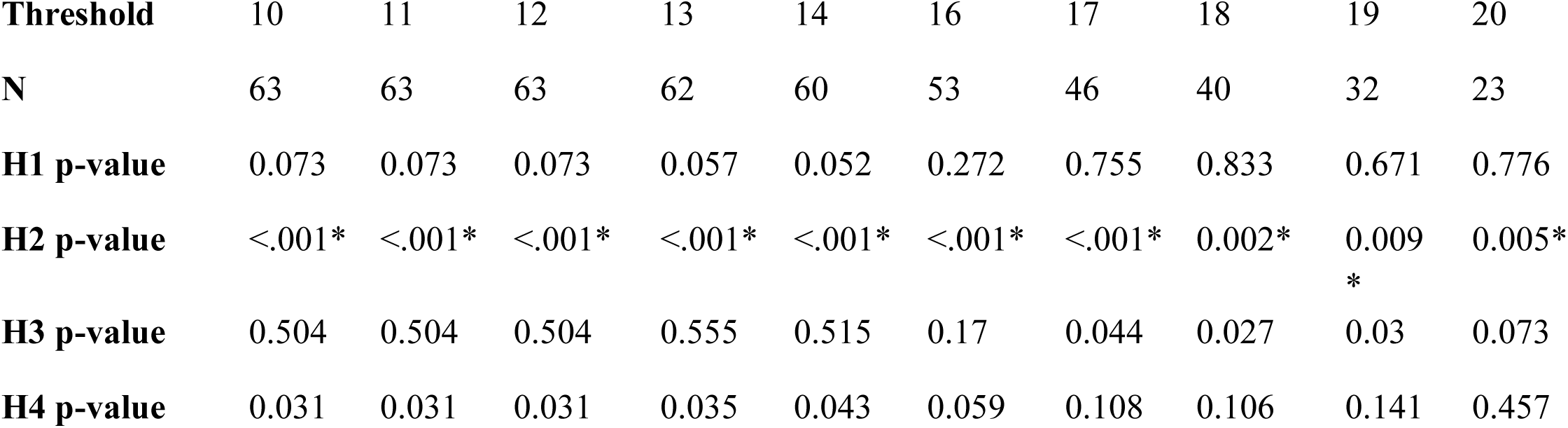
Number of included participants and corresponding p-values across different minimum trial thresholds per condition and weight.

## Further Exploratory Analyses

### The relationship between MEP and perceptual differentiation between heavy and light lifts

We tested whether the difference between normalised peak-to-peak MEP amplitude during the observation of light and heavy lifts would correlate with the difference in subjective heaviness ratings in the cued and unpredictable conditions. While the relationships were not significant, in neither the unpredictable condition, r = -0.15, p = 0.28, nor the cued condition, r = 0.00079, p =1, the tendency was consistent with our results, suggesting that, in the unpredictable condition, participants who showed greater difference between heavy and light objects in their MEP amplitudes may also show greater differentiation in perceived heaviness. This was, however, not statistically supported and should be interpreted as a hypothesis at most.

**Figure 6.**
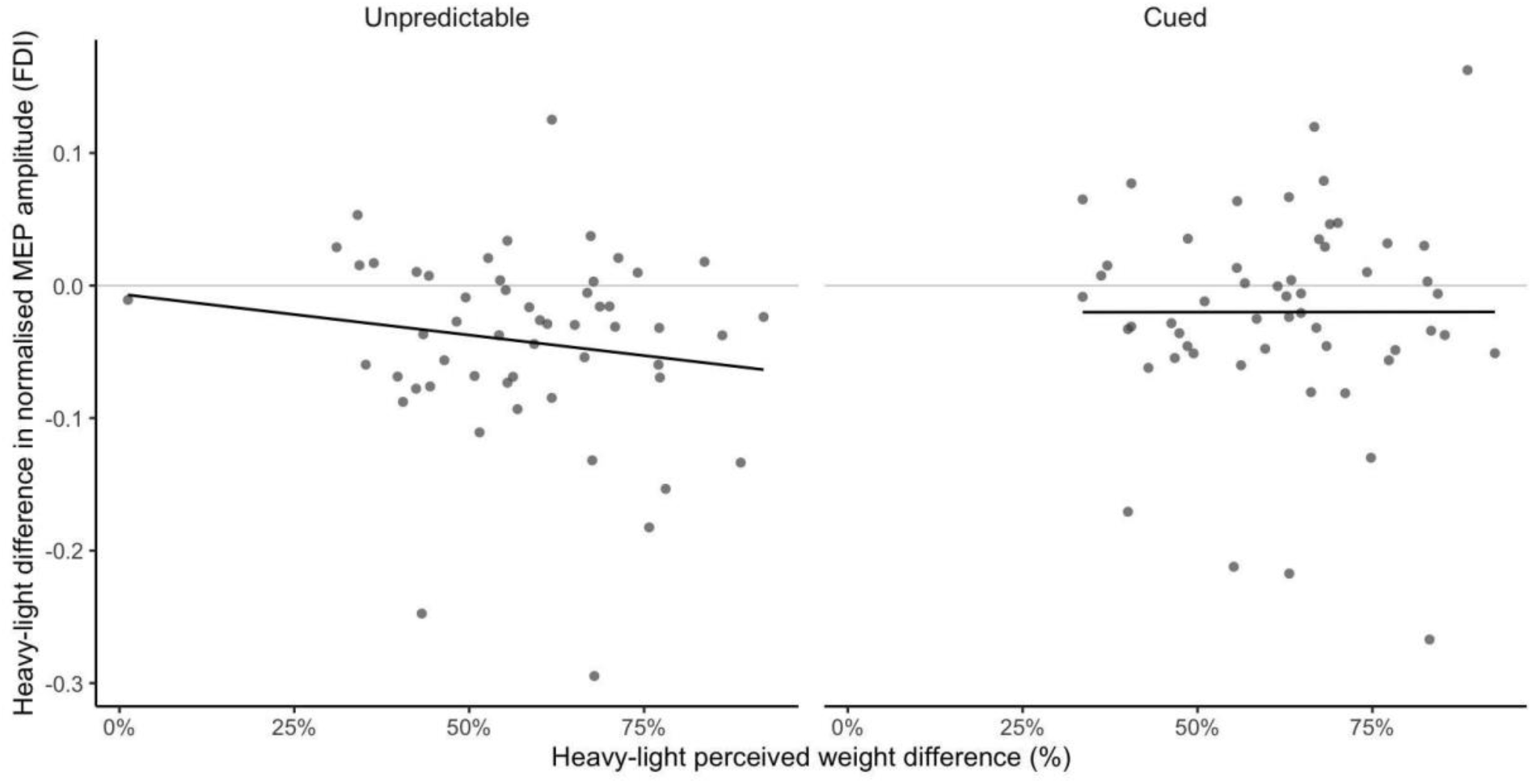
Relationship between weight-related changes in MEP amplitude and perceived heaviness. Dots show each participant’s heavy-light difference in mean normalised FDI MEP amplitude plotted against the same participant’s heavy-light difference in perceived heaviness ratings, separately for each condition. Note, heaviness ratings were collected separately in the weight-judgement task after the TMS experiment.

### Analyses on APB muscle instead of FDI

We repeated the four preregistered analyses looking at the normalised MEP peak-to-peak amplitude in the APB instead of the FDI muscle to assess whether the findings were specific to the FDI. Note that, based on the same inclusion criteria as for FDI, data from only 51/57 participants were available for this analysis. This was because, during data collection, noise reduction was optimised for the channel recording from the FDI muscle; for example, the experiment only started if the noise level was acceptable, but this was less strictly applied to the channel recording from the APB muscle. In the APB muscle, MEP responses were larger during the observation of light than heavy lifts in all three conditions: unpredictable, blocked, and cued. The difference between the blocked and unpredictable conditions (H3) was not significant, suggesting that the size of this difference did not differ between conditions. The full statistics for the APB analyses are summarised in Table 4 and visualised in Figure 7.

**Figure 7.**
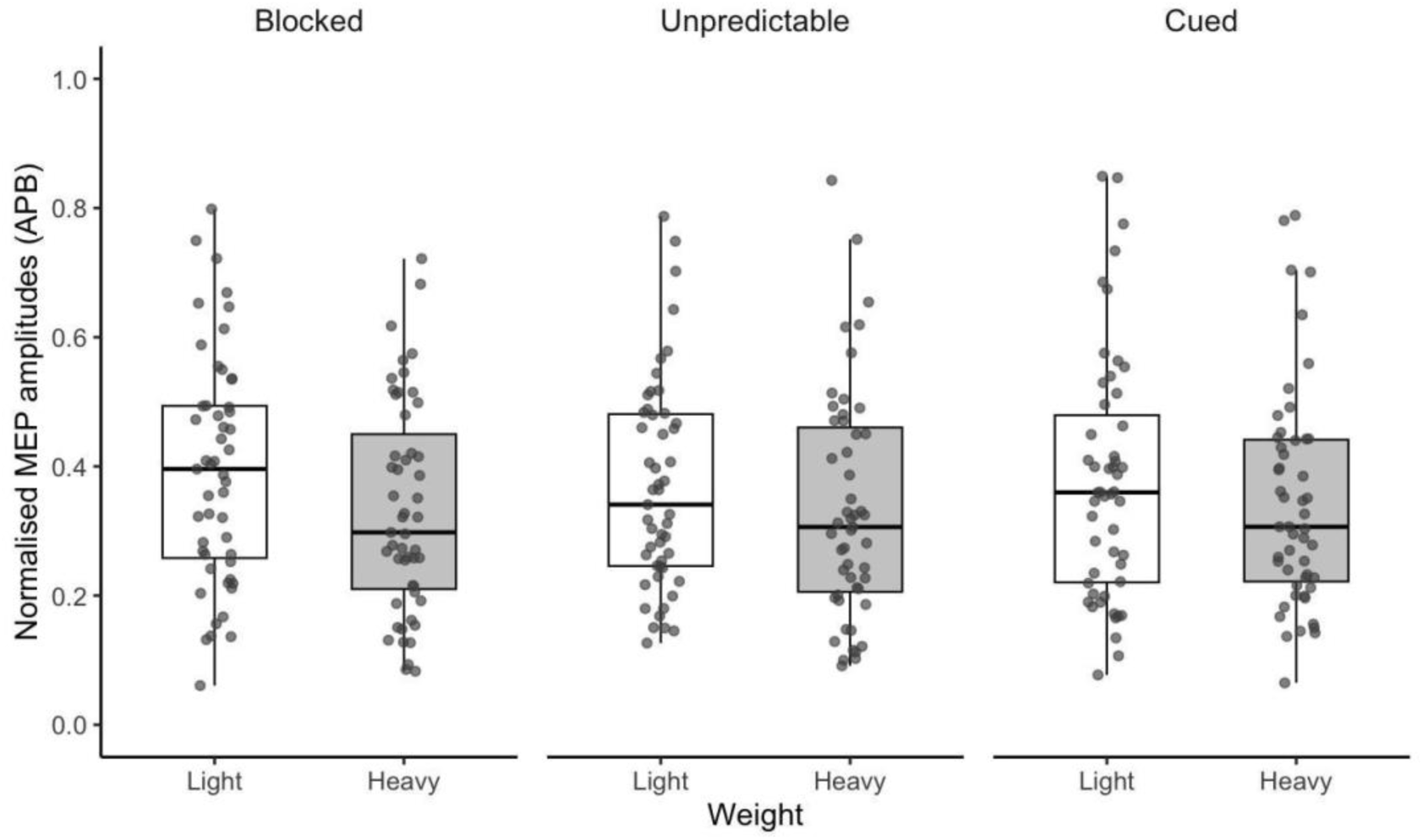
Normalised peak-to-peak MEP amplitudes for light and heavy object lifts in the blocked, unpredictable, and cued conditions in the APB muscle. Points show participant-level condition means (n=51), the boxes show the interquartile range, whiskers show 1.5 x the interquartile range, and the horizontal lines show medians.

**Table 4.** Results of the exploratory APB analyses.

| Analyses | N | Mean<br>diff. | 95%<br>CI<br>lower<br>bound | 95% CI<br>upper<br>bound | t-value | df | p-value | dz | BF10 | BF01 |
| --- | --- | --- | --- | --- | --- | --- | --- | --- | --- | --- |
| H1 | 51 | -0.06 | -0.091 | -0.029 | -3.88 | 50 | < .001* | -0.54 | 81.26 | 0.01 |
| H2 | 51 | -0.034 | -0.056 | -0.012 | -3.11 | 50 | 0.003* | -0.44 | 10.46 | 0.1 |
| H3 | 51 | -0.026 | -0.06 | 0.007 | -1.59 | 50 | 0.119 | -0.22 | 0.49 | 2.03 |
| H4 | 51 | -0.034 | -0.057 | -0.01 | -2.81 | 50 | 0.007* | -0.39 | 5.08 | 0.2 |

## Discussion

We investigated the effect of trial predictability on the previously reported effect of larger MEP responses during the observation of heavy compared to light lifts. Although the present study was motivated by the possibility that predictive cues in previous studies (Alaerts, Senot, et al., 2010; Alaerts, Swinnen, et al., 2010) contributed to this effect, our findings did not support the idea that predictability alone is sufficient to produce it. On the contrary, we found that only in the condition with no predictive cues (neither visual nor trial-presentation), the corticospinal system responded more to TMS during the observation of light than of heavy weights. This effect was robust across multiple analyses. The same effect was found when repeating the analysis using raw MEP peak-to-peak amplitude rather than normalising to the largest MEP per participant, when using an alternative outcome measure of AUC instead of peak-to-peak amplitude, and when examining responses in the APB muscle rather than FDI. In the APB, however, larger MEPs during observation of light than heavy lifts were present in all three conditions, suggesting that the larger response during light lifts is not muscle-specific, whereas the influence of trial predictability may be. We tested these effects on a much larger sample than previous studies, in a Registered Report, and have confirmed that results were robust to any deviations from the initial protocol.

The evidence for this effect (H2) was very strong, with the data being 223.64 times more likely under the alternative than under the null hypothesis and the size of this effect, dz = 0.56, exceeded the threshold we preregistered as the minimum effect size with theoretical relevance (dz=0.52). However, it reflected only an approximately 4% change in normalised peak-to-peak MEP amplitude. Moreover, the direct comparison between the blocked and unpredictable condition (H3) did not confirm that the MEP response during the observation of light and heavy weights differed significantly across conditions. Given that this finding is in the opposite direction to what was hypothesised, it warrants direct replication before firm conclusions can be drawn.

We initially included the cued condition (where the heavy and the light objects were different sizes) as a positive control, expecting that the top-down prediction coming from the visual cues would enhance MEP differences between heavy and light lifts beyond that driven by kinematics alone. The lack of effect in this condition is inconsistent with previous studies showing that visual cues about object weight can modulate corticospinal excitability and suggests that the effect of predictive cues might be heavily context dependent. The visual distinction between the size of our 200 g and 1800 g stimuli was likely weaker than in previous studies where object size or shape may have provided more salient cues. For example, Alaerts et al. (2010) used objects that were visually distinct in size, shape and function. Similarly, Buckingham et al. (2014) found size-based modulation of corticospinal excitability when participants observed lifts of identically weighted cubes, but the size contrast in that study was more pronounced than in the present study (Figure 1). Another consideration is that, although pulse timing was matched between heavy and light lifts relative to object contact, the cued condition was the only condition not matched in absolute timing from video onset and this may have impacted the results. In this condition, object contact occurred earlier relative to absolute timing from video onset for the small-light object (2.3 seconds) than for the large-heavy object (3 seconds). This is not a deviation from the preregistration, but a post-hoc observation about the nature of the natural lifting videos with an actor who is unaware of the weights. We elaborate on this below.

### The critical problems of choosing a reference point for stimulation timing during the observation of heavy and light lifts

The choice of reference point on how stimulation timing is synchronised across conditions might explain some of the inconsistencies in the literature. During the observation of heavy and light lifts, stimulation could be matched based on at least three different reference points: time of contact with the object, phase of the movement, or (while presumably less important) the start of the video. Each of these reference points has different trade-offs. In *natural* lifts, synchronising stimulation timing based on any of these almost guarantees a mismatch on at least one other, because heavy and light lifts naturally unfold with different speeds. As corticospinal excitability can change over the course of a trial due to temporal preparation/expectancy (Hasbroucq et al., 1997) any of the three mismatches could influence MEP amplitudes independently of object weight.

In the studies by Alaerts et al., stimulation timings were based on when the velocity of the vertical displacement of the lifted object exceeded 20 mm/s. This synchronises the stimulation in terms of the dynamics of the movements, but because this velocity is reached later for the heavy lifts (see Figure 1 in Alaerts et al 2010 or Alaerts et al 2012), stimulation in the heavy condition occurs systematically later relative to object contact compared to the light condition. In contrast, in our study and in the study by Senot et al. (2011), pulses were delivered within a fixed window after contact with the object for both light and heavy lifts. The limitation of matching based on contact is, however, that it does not guarantee that the videos (or observed live action in the case of Senot et al.) will be matched for the phase of the lift. If heavy objects have a longer loading phase (again, see Figure 1 in Alaerts et al 2010 or Alaerts et al 2012), then a larger proportion of a fixed stimulation window might include grasp for the heavy than for the light lifts.

Lastly, there might also be an anticipatory effect based on the time from the start of the video, although this is likely the least important concern out of the three reference points, especially when stimulation is randomised within a video and the inter-stimulus interval also varies, as in our study. This was not a concern in the blocked and unpredictable conditions in our study. Each video started at the same time as the reach, and because in these conditions the heavy and light objects looked identical, and the actor was unaware of the weights, there was no difference in how he reached for them (to the level that could not be compensated by a few hundred milliseconds of editing). The touch of the object was at the same time, 3 seconds after the start of the video (and therefore also 3 seconds after the start of the reach) so stimulation timing was matched both for video onset and object contact. However, this was only possible in these conditions because the objects looked the same, and could not be achieved in the cued condition.

There is no simple solution for these trade-offs for the different timing reference points in naturalistic lifting. Perfectly timed or edited videos would improve experimental control, but would remove some of the natural information from kinematics/dynamics, which would make the paradigm less ecologically valid. Very importantly, these timing trade-offs cannot explain the difference between the blocked and unpredictable conditions in our study, which used the exact same heavy and light videos with the same timing window, but they may explain inconsistencies between our findings and studies using different stimulus sets and timing choices.

## Limitations

One limitation of the present study, as in many action-observation TMS studies, is that we have limited direct information about the cognitive processes occurring while participants observed the lifts. For example, we cannot know on each trial whether participants were attending to the weight of the object, the kinematics of the lift, or other aspects of the stimulus. Some previous studies have included additional attention checks during the task, such as asking participants to judge the timing of a sound relative to the observed action (Senot et al., 2011). However, such tasks may also change how participants observe the action and still do not provide a direct measure of weight processing. In the present study, we tried to reduce this limitation by including regular breaks after every 11 trials and by asking participants to complete weight judgements after the TMS task. These behavioural judgements confirmed that participants could reliably distinguish between the light and heavy lifts.

A second limitation concerns the stimulation-timing error that led to the protocol deviations from the Stage 1 manuscript. We found no evidence that this timing error introduced bias into the results. However, the experiment was not designed to test whether the timing error had broader effects on attention, temporal expectations, or stimulus predictability. Such effects are unlikely to have changed the conclusions, because the pattern of results was unchanged when the longer-timing trials were included or excluded. In the cued condition, results were not meaningfully altered by retaining or excluding the additional 0.5 s trials, and the blocked-condition order comparison did not reach the preregistered significance threshold. Nevertheless, it remains theoretically possible that the additional 0.1 s in the longer stimulation window affected corticospinal responses, but this was likely in a way that did not impact the results.

## Implications

While this study focused on the effect of trial predictability on corticospinal excitability during the observation of heavy and light lifts in neurotypical individuals without movement problems, it led us to generate some hypotheses about potential implications. First, the results suggest that observing more forceful movements does not necessarily mean the observer’s corticospinal system is more strongly engaged, and consequently that such observation would not be more beneficial for motor learning/rehabilitation. Secondly, they raise the possibility that the predictive trial presentation may negatively influence the corticospinal system’s ability to differentiate between movements. These implications may be relevant for observation-based physiotherapeutic rehabilitation (e.g., see Buccino 2014; Giannakopoulos et al., 2022), but should be tested directly in those relevant contexts.

## Conclusion

Our findings provide no evidence that the corticospinal system responds more to TMS during the observation of someone lifting a heavy object compared to a light one. In the FDI muscle, the only reliable differentiation between heavy and light lifts occurred when predictive information was unavailable. This suggests that, in the absence of predictable information, the corticospinal system was sensitive to the observed kinematics, but, contrary to previous findings, its excitability varied inversely with object weight.

